# Cryo-EM structures of apo human Factor XIa reveal catalytic-domain flexibility and exposure of the Factor IX-binding site

**DOI:** 10.64898/2026.08.25.744604

**Authors:** Alena I. Siutkina, Alexander Neuhaus, Marvin Taterra, Marcel Bermúdez, Christos Gatsogiannis, Dmitrii V. Kalinin

## Abstract

Factor XI (FXI) is a key coagulation protease of the intrinsic pathway of blood coagulation and an emerging antithrombotic target. However, the structural transition from zymogen to active Factor XIa (FXIa) has remained poorly understood. Using cryo-EM, we demonstrate that FXI activation results in a global reorganization of the homodimer, extending beyond the activation loop to include a significant reorientation of the catalytic domain (CD) relative to the apple-domain (AD) platform. The CD displays pronounced conformational heterogeneity; we identify three distinct conformers, suggesting that FXIa exists as a dynamic ensemble rather than a single rigid state. MD analysis indicates that activation disrupts the inter-CD allosteric communication present in the zymogen, thereby facilitating this flexibility. CD plasticity allows for the dynamic exposure of the A3 exosite, facilitating the binding of Factor IX. Comparison with plasma kallikrein (PKa) suggests that such structural flexibility may be a shared feature of apple-domain-containing contact-system proteases. Our results reveal that FXIa functions as a dynamic ensemble, providing a structural framework for understanding substrate recognition and identifying novel, non-catalytic sites for the development of specific FXIa inhibitors.

**Key Points:**

- First apo cryo-EM structure of full-length FXIa reveals the unliganded active architecture and exposes the cryptic A3-domain FIX-binding exosite.
- Three distinct apo-FXIa conformations and MD simulations reveal catalytic-domain flexibility during the transition from zymogen-like FXI to active FXIa.

## Introduction

Coagulation factor XI (FXI) is a structurally and functionally unique component of the intrinsic pathway of blood coagulation. Unlike most coagulation protease zymogens, FXI circulates as a disulfide-linked homodimer, with each subunit composed of four N-terminal apple domains (ADs) (A1-A4) and a C-terminal trypsin-like catalytic domain (CD).^1–3^ The ADs form a planar disk-like platform, with A1 and A2 arranged antiparallel to A3 and A4, whereas the CD rests on this platform in a characteristic “cup-and-saucer” architecture.^1,3,4^ In the dimer, the two AD disks are inclined relative to each other, generating a V-shaped arrangement stabilized by the A4 domains, including the intersubunit C321-C321 disulfide bond and additional hydrophobic and electrostatic interactions. Upon proteolytic cleavage after R369, each FXI subunit is converted into activated FXI (FXIa), in which the apple-domain-containing heavy chain remains linked to the catalytic light chain by a disulfide bond (Figure 1A).^1–3,5^ FXIa amplifies thrombin generation primarily through activation of factor IX (FIX), thereby supporting thrombus propagation and clot stability.^2,6^

**Figure 1.**
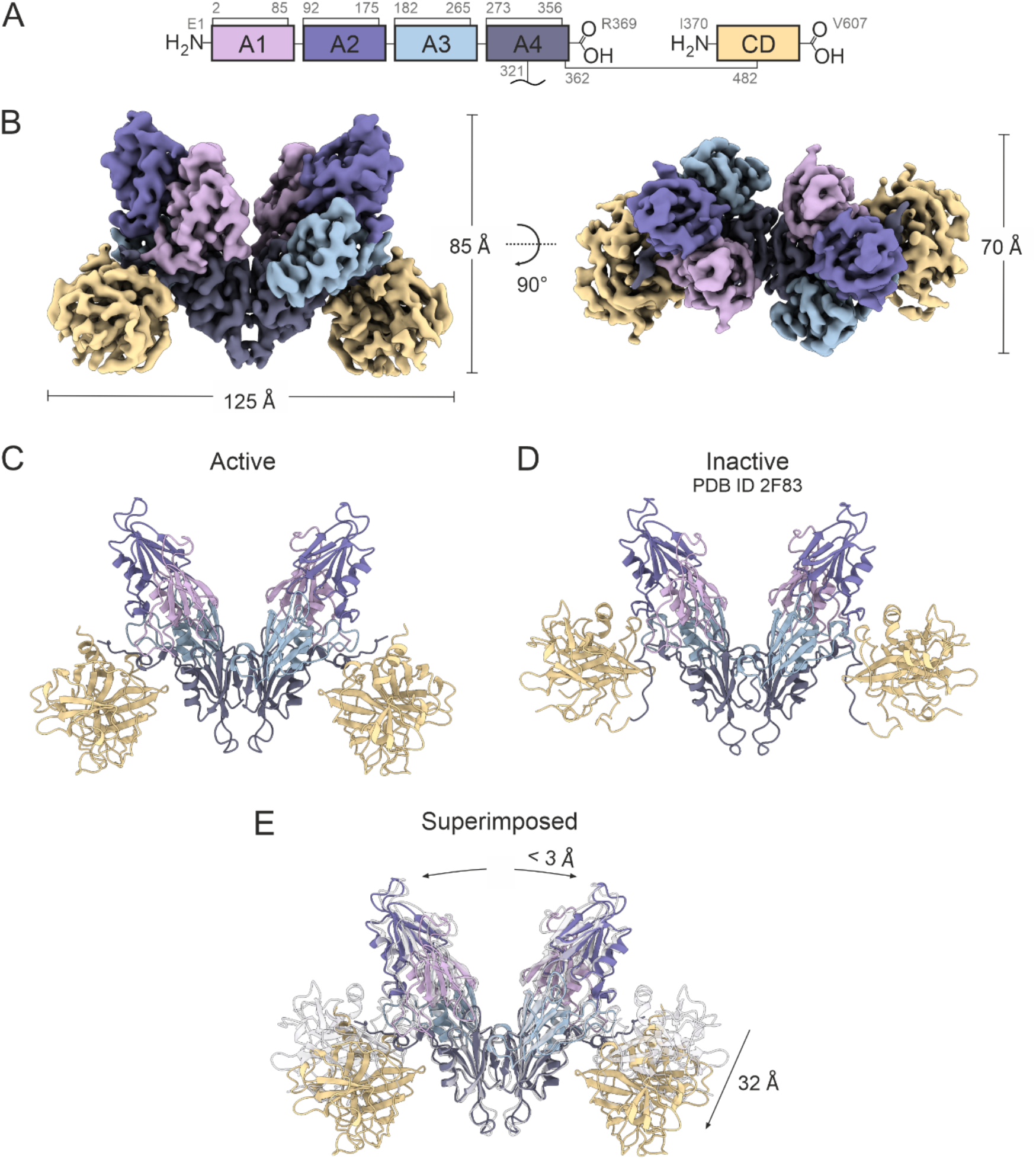
Cryo-EM structure of human FXIa. (A) Schematic representation of the domain architecture of the FXIa homodimer. The disulfide bonds involved in the enclosure of apple domains or connection with other domains are shown in grey together with the corresponding residue numbers. (B) Cryo-EM reconstruction of the FXIa homodimer at 2.80 Å resolution is shown from two perspectives. Domain colors correspond to those shown in panel A. (C) Atomic model of the FXIa homodimer built into the cryo-EM reconstruction. (D) Previously reported structure of the FXI in its inactive (zymogen) state (PDB ID: 2F83^3^). (E) Superimposition of the present FXIa homodimer structure with the previously reported structure of the FXI zymogen^3^ illustrates the major differences between the two states.

FXI has emerged as an attractive anticoagulant target because it appears to contribute more strongly to pathological thrombosis than to normal hemostasis.^7^ Individuals with congenital FXI deficiency, historically referred to as hemophilia C, typically show variable and often mild trauma- or surgery-associated bleeding. At the same time, epidemiological and clinical data indicate reduced thrombotic risk in individuals with low FXI levels.^2,8^ This separation between thrombosis and hemostasis has stimulated major pharmaceutical interest in FXI/FXIa inhibition, including antisense oligonucleotides, monoclonal antibodies, and small-molecule FXIa inhibitors such as milvexian and asundexian, several of which have advanced into clinical development.^8–11^

Beyond its role in FIX activation, FXI functions as a multi-domain scaffold for several biologically important interactions. The A1 domain contributes to thrombin and high-molecular-weight kininogen binding; the A2 domain participates in high-molecular-weight kininogen binding and FXIIa-dependent mechanisms; the A3 domain mediates interactions with FIX, platelet GPIbα, polyphosphate, and other ligands; whereas the A4 domain is central to dimerization and contributes to FXIIa- and thrombin-dependent activation.^2,8^ Thus, defining the architecture of full-length FXIa is important not only for understanding FIX recognition but also for placing these domain-specific interaction surfaces in the context of the activated enzyme.

Despite this biological and therapeutic importance, the structural basis of FXI activation and substrate recognition has remained incompletely understood. The crystal structure of the FXI zymogen provided a detailed view of the compact “cup-and-saucer” architecture (Figure 1D).^1,3,4^ Previous low-resolution electron microscopy and small-angle X-ray scattering studies suggested that conversion of FXI to FXIa is accompanied by large conformational changes.^12^ In line with this, FIX does not bind the FXI zymogen, indicating that activation must create or expose a substrate-recognition surface that is not available in the inactive state.^2,6^ Biochemical and mutagenesis studies previously implicated the A3 domain, particularly residues around I183, R184, D185, and the C-terminal region of A3, in binding the FIX Gla domain.^6^ In the zymogen structure, this proposed FIX-binding exosite is masked by the CD, suggesting that FXI activation requires a major repositioning of the CD to expose a cryptic A3-domain surface required for FIX engagement.^2,6^

Several crystal structures of the FXI zymogen,^3,4,13^ individual FXI domains,^4,12,14^ and the isolated activated catalytic domain^15–35^ have been reported. While our work was being completed, cryo-EM structures of FXIa in complex with two mechanistically distinct antibody fragments, amrecibart and cenvacibart, were reported.^36^ These antibody-bound structures provided the first reported structural description of full-length FXIa and revealed activation-associated displacement of the CD relative to the AD platform, together with exposure of the A3-domain surface.^36^ While these structures offered valuable insights into the activated FXIa architecture, the primary focus of that study was the biological and anticoagulant characterization of amrecibart and cenvacibart.^36^ The structural analysis mainly rationalized their domain-specific modes of FXI/FXIa inhibition and did not study deeply the activation mechanism. The unliganded architecture of FXIa and the detailed surface organization of apo-FXIa remained to be defined.

Here, we report cryo-EM structures of full-length human apo-FXIa, providing, to our knowledge, the first structural description of activated FXIa in the absence of bound antibodies, inhibitors, or other ligands. At 2.8 Å resolution, the main apo-FXIa reconstruction was obtained at a higher resolution than the recently reported antibody-bound FXIa structures (3.2 Å)^36^, allowing for the visualization of the activated enzyme in previously unavailable detail. Comparison with zymogen FXI and antibody-bound FXIa confirms that activation is accompanied by a global architectural rearrangement, with some changes across the AD platform and pronounced repositioning of the CD. This movement uncovers a cryptic surface on the A3 domain, providing a structural basis for FIX binding. Molecular docking supports the formation of a FIX-binding interface at this newly exposed A3 pocket, while molecular dynamics simulations provide a dynamic view of the transition from the FXI zymogen-like arrangement toward the activated FXIa state. Importantly, our cryo-EM analysis reveals pronounced conformational heterogeneity of the CD, with several apo-FXIa conformations observed in the absence of bound ligands. Together with molecular dynamics simulations, these findings show that activated FXIa is not a single rigid structural state but a dynamic ensemble of catalytically competent conformations. These insights provide a mechanistic foundation for a deeper understanding of FXIa physiology and the design of more effective and safe modulators.

## Methods

### FXIa protein

Purified human coagulation factor XIa (FXIa) was purchased from Innovative Research (Product SKU: IHUFXIA100UG; lot no. 6112). According to the certificate of analysis, FXIa was prepared from human plasma-derived FXI by activation with human FXIIa, followed by removal of FXIIa using a corn trypsin inhibitor affinity column. The protein was supplied as a frozen liquid in 4 mM sodium acetate, 0.15 M NaCl, pH 5.3, and stored at −80 °C until use. The supplied preparation had a concentration of 1.27 mg/mL, a volume of 0.079 mL, and a reported activity of 223.9 IU/mg. The molecular weight was reported as approximately 160 kDa under non-reducing SDS-PAGE conditions, and the purity was >95% as determined by SDS-PAGE analysis. The protein was used for negative stain and cryo-EM sample preparation without further purification.

### Negative stain EM analysis

For negative stain EM sample preparation, 5 µL of the protein sample, diluted to a concentration of 0.008 mg/mL, was applied to a glow-discharged carbon-coated copper grid and incubated for 1 min at room temperature. The excess protein solution was removed by blotting with Whatman (grade 4–5) filter paper, followed by washing with 2 × 10 μL deionized water and 2 × 10 µL 0.75% uranylformate solution. The final uranylformate droplet was incubated for 75 s before blotting and air-drying the grid.

Image acquisition was performed using a Talos L120C G2 TEM operating at an acceleration voltage of 120 kV. Datasets were acquired with a 4k × 4k CETA-F scintillator camera at a defocus of −1 μm and a pixel size of 1.2 Å/px.

### Sample vitrification and cryo-EM data acquisition

For preparing the grids for cryo-EM, two samples of 5 μL of FXIa at concentrations of 0.4 mg/mL and 0.1 mg/mL were applied onto freshly glow-discharged UltrAuFoil R1.2/1.3 holey gold grids (Quantifoil). After excess liquid was removed, the samples were vitrified in liquid ethane using a Vitrobot II automatic plunge-freezer (Thermo Fisher Scientific).

### CryoEM data processing and 3D reconstruction

Datasets were acquired with a 300 kV Titan Krios G4 microscope (Thermo Fisher Scientific) equipped with an E-CFEG, a Selectris X energy filter, and a Falcon 4i direct electron detector operated by the software EPU (Thermo Fisher Scientific). A total of 18179 (0.4 mg/mL) and 44.805 (0.1 mg/mL) micrographs were collected in electron event representation mode (EER) at a nominal magnification of 215k, corresponding to a pixel size of 0.571 Å/px. The majority of micrographs were collected at a defocus of −0.8 to −2.8 μm. The Selectris X energy filter was used for zero-loss filtration with a slit width of 10 eV. A total dose of ~50 e−/Å^2^ (estimated shortly after freshly flashing the cold-FEG) was set by adjusting the exposure time to 2.05 s per micrograph.

### Image processing and 3D reconstruction

Data processing was performed in CryoSPARC^37^ (Supplementary Figure 1), using C2 symmetry.

### Model building and validation

The model was manually built in ChimeraX v1.9^38–40^ using the Isolde plugin.^41^ The initial model for the structure was the published zymogen structure with PDB ID 2F83^3^ and 5I58^4^. The resulting model was further refined using a combination of Isolde and Phenix. Multiple rounds of the above adjustments were performed until the model sufficiently described the experimental map.

### Visualization and analysis of cryoEM maps and models

Visualization, analysis, and figure preparation were done with ChimeraX v1.9^38–40^ using the Isolde plugin^41^ and CorelDRAW (https://www.coreldraw.com). Local resolution gradients within a map were calculated with CryoSPARC^37^ and visualized with ChimeraX v1.9.^38–40^

### Molecular docking

In addition to the newly reported structures, the zymogen structure (PDB: 2F83^3^) was retrieved from the Protein Data Bank and prepared using *MOE 2024.0601* (Molecular Operating Environment, Chemical Computing Group ULC). For the zymogen, missing loop regions were remodeled, glycosylation was added, and connectivity was adjusted to match the active state structure. Truncated ends were patched, missing atoms were added, and steric clashes as well as Ramachandran outliers were minimized using the AMBER14:ETH force field. All systems were protonated at pH 7.4. Protein-protein docking was performed in *MOE 2024.0601* using the prepared active-state structure and the Gla domain of FIX (PDB: 1NL0^42^) using default settings. Rigid-body docking was employed to preserve the initial geometry of the ω-loop, which is natively coordinated by calcium ions. The resulting poses were subsequently subjected to visual inspection, including assessment of Ramachandran outliers and steric clashes. Clashes arising from the rigid-body docking were relieved by energy minimization to an RMS gradient of 1×10^−5^ kcal mol^−1^ Å^−1^ until no clashes remained. Ligandscout^43^ was used to analyze the resulting protein-ligand interactions, and ChimeraX v1.9^38–40^ was used for visualization.

### Molecular dynamics simulations

Simulation systems were assembled in CHARMM-GUI^44^ and solvated in a cubic TIP3P water box containing 0.15 M NaCl with an enlarged padding of 20 Å. Simulations were performed with OpenMM 8.4^45^ using the CHARMM36m force field^46^ on NVIDIA RTX 4090 GPUs (NVIDIA Corporation, Santa Clara), following the CHARMM-GUI minimization and equilibration protocol^44^. Throughout, bonds involving hydrogen were constrained and rigid water geometry was enforced with SETTLE. Nonbonded interactions used a 1.2 nm cutoff, with a CHARMM force-switch applied to van der Waals interactions between 1.0 and 1.2 nm and PME electrostatics computed with a 1.2 nm real-space cutoff. The system was first energy-minimized with OpenMM’s L-BFGS local minimizer (maximum 5000 iterations, or until the root-mean-square force fell below 100 kJ mol^−1^ nm^−2^), then equilibrated for 125 ps at a 1 fs timestep in the NVT ensemble. Initial velocities were drawn from a Maxwell–Boltzmann distribution at 303.15 K, and the temperature was maintained by a Langevin integrator (friction coefficient 1 ps^−1^) with no pressure coupling. During both stages, flat harmonic positional restraints were applied to the protein backbone (400 kJ mol^−1^ nm^−2^) and side-chain heavy atoms (40 kJ mol^−1^ nm^−2^), together with dihedral restraints on the carbohydrate (glycan) atoms (4.0 kJ mol^−1^ rad^−1^), relative to their initial coordinates. Production molecular dynamic (MD) simulations were continued from the final equilibrated configuration of each system with all restraints removed and propagated in the NPT ensemble with a 2 fs timestep. The temperature was held at 303.15 K by the Langevin integrator and the pressure at 1.0 bar by an isotropic Monte Carlo barostat with volume-move attempts every 100 steps. Each system was simulated in quintuplicate, with a 500 ns trajectory and 5000 frames recorded per replica. These unbiased ensembles were subsequently used for enhanced sampling of the state transitions, and the parametrized molecular mechanical (MM) structures served as the basis for quantum mechanical/molecular mechanical QM/MM calculations (full details of both are provided in the Supplementary Methods). Long-range allosteric communication was analysed with MDPath^47^ using default settings. Replicas were pooled and evaluated with confidence analysis (https://github.com/wolberlab/mdpath/pull/110).

## Results

### Cryo-EM structure of human FXIa

Plasma-derived FXIa was subjected to cryo-EM, yielding a structure at an average resolution of 2.8 Å with C2 symmetry imposed (Figure 1A, B). Consistent with previously reported data, each monomer comprises four ADs, A1-A4, which form a saucer-shaped AD disc, and one cup-shaped CD.^4,36^ Superimposition of the FXIa homodimer model (Figure 1C) with the FXI zymogen structure (PDB ID: 2F83^3^; Figure 1D) reveals a 32 Å displacement of the CD, measured at G533, as well as a movement of up to 3 Å of each AD disc (Figure 1E).

### Structural rearrangements accompanying FXI activation and potential basis for FIX recognition

Activation of each FXI subunit is initiated by proteolytic cleavage of the peptide bond between R369 and I370 (activation loop), generating I370 as the new N-terminus of the CD.^48^ Comparison of the zymogen with the active FXIa structure reveals that this cleavage is accompanied by extensive conformational rearrangements: both parts of the activation loop are repositioned in opposite directions, while the entire CD undergoes a rotation and a displacement relative to the AD disc (Figure 2A). In the zymogen, the activation loop contributes to the electrostatic landscape surrounding the CD (Figure 2B, upper left). After cleavage, the D359–R369 segment undergoes a dramatic rotation towards the opposite side of the CD (Figure 2B, lower right). Comparison of the interactions between CD and the D359– R369 fragment in both states reveals an increase in the number of non-covalent contacts (Supplementary Figure 2), which can explain the enhanced stabilization within the newly occupied groove. In addition, the movement of the CD exposes the previously hindered I183– R184–D185 motif on the A3 domain, which has been previously implicated in FIX activation (Figure 2B)^6^. Next, we asked which structural changes accompany or result from cleavage beyond static structures. We therefore simulated the zymogen-like to active-like transition using enhanced sampling and extracted the slowest concerted motions with DeepTICA.^49^ Cleavage was accompanied by reciprocal rearrangements at the R369–I370 site: I370 docked into the activation pocket and formed the I370–D556 salt bridge (Supplementary Fig. 3a, left), whereas R369 moved to the opposite face of the CD, where an E361–R479 salt bridge stabilized the liberated heavy-chain latch (Supplementary Fig. 3b). Both rearrangements varied little along the transition, identifying them as immediate consequences of cleavage. The dominant slow motion instead involved the CD core itself, which consolidated gradually around the coevolving A375–Q515 contact (Supplementary Fig. 3a, middle), followed by formation of the S376–L514 hydrogen bond (Supplementary Fig. 3a, right). Together, these results support a two-step activation model in which cleavage sets the activation pocket and releases the latch immediately, whereas slower consolidation of the core stabilizes the mature active conformation. QM/MM calculations indicated that forming the activation-induced I370– D556 salt bridge results in a more negative potential for the catalytic triad (Supplementary Fig. 3c), which drives preorganisation of the active site for catalysis.

**Figure 2.**
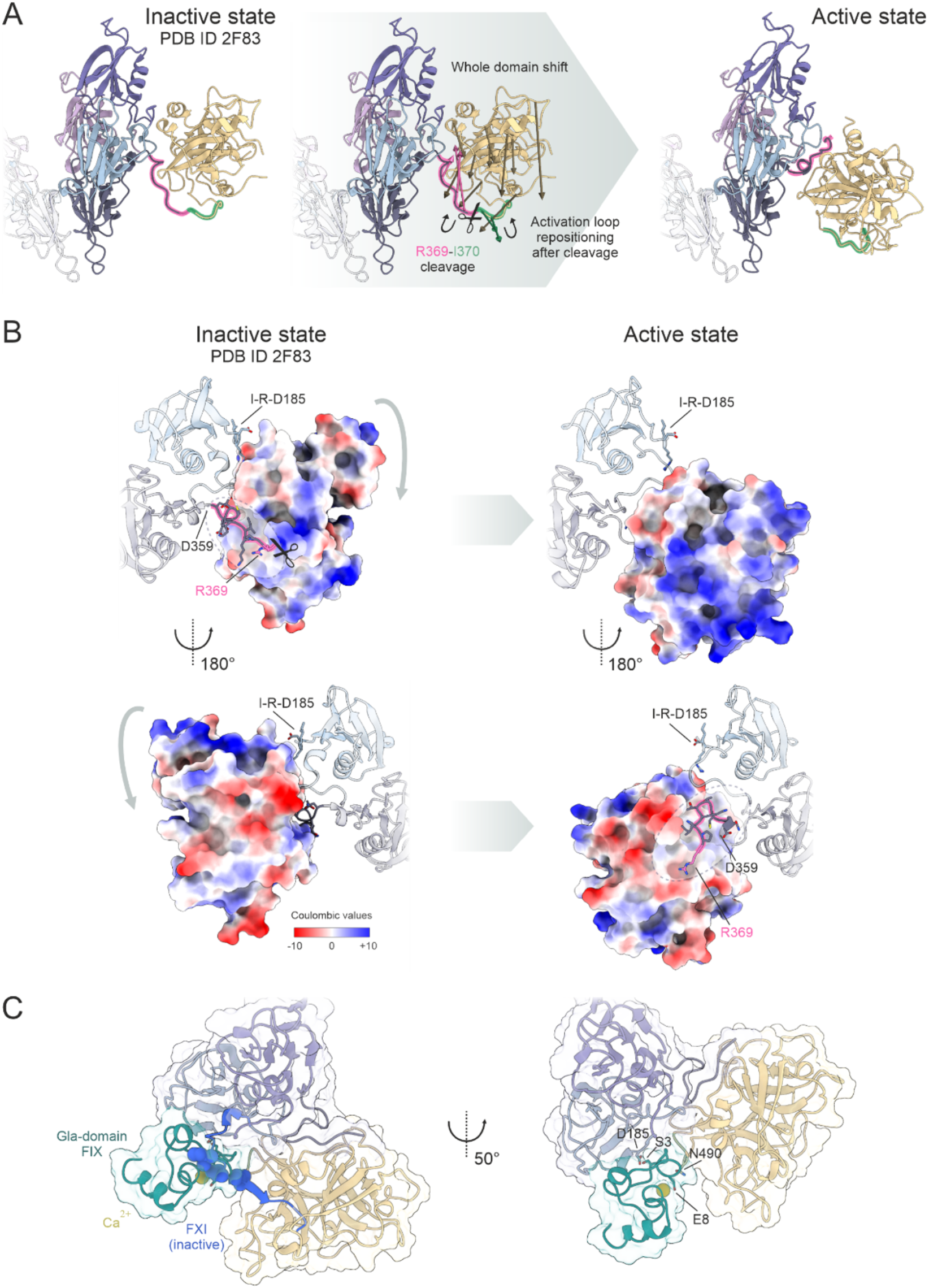
Structural rearrangements accompanying FXI activation and potential basis for FIX recognition. (A) Comparison of the inactive FXI zymogen (left, PDB ID: 2F83^3^) and the active FXIa structure (right). The central schematic illustrates the structural transition following proteolytic cleavage at the R369–I370 activation site. Residues D359–R369 are highlighted in pink, while residues I370–R378 are shown in green. Cleavage induces repositioning of the activation loop together with a rotation and displacement of the CD, as indicated by the arrows. (B) Electrostatic surface representation of the CD before and after activation. The orientation of the newly generated N-terminus (residues D359–R369) and the conserved I183–R184–D185 (IRD) motif are shown relative to the CD surface. Two views, rotated by 180°, illustrate how activation alters the electrostatic landscape and the accessibility of these structural elements. (C) Overlay of inactive FXI (PDB ID: 2F83^3^) and active FXIa on the protein–protein docking model of the FIX Gla domain (PDB ID:1NL0^42^, green). In the inactive state, before cleavage, the K486–D495 loop (dark blue) fully occludes the Gla-domain binding site, preventing engagement of FIX by inactive FXI. Residues directly competing with the Gla-domain are displayed as blue spheres.

To explore whether this rearrangement could facilitate FIX recognition, the FIX Gla domain, which has previously been implicated in the interaction with FXIa,^50^ was docked into both the inactive FXI and the activated FXIa structures (Figure 2C). In the inactive conformation, docking did not result in engagement of the FIX Gla domain by the FXI pocket. In contrast, the activated structure presents a sterically favorable interface that accommodates the FIX Gla domain. In the Ca^2+^-coordinated ω-loop conformation of the Gla domain, the carboxylate groups are internalized, exposing a predominantly hydrophobic surface. In the docked complex, this hydrophobic surface inserts into a hydrophobic pocket on FXIa. Only limited direct polar contacts are observed, involving S3 and E8 of the Gla domain, indicating an interaction driven largely by hydrophobic contacts rather than specific polar interactions. Given that this pocket on the A3 domain of FXIa is hydrophobic and highly solvent-exposed, displacement of water molecules from it likely constitutes a major energetic determinant of binding.

A closer comparison of the FXI zymogen and FXIa revealed additional substantial rearrangements within the AD discs and at the CD-apple-domain interfaces. In the inactive state, the CD occupies a cavity between the A2 and A3 domains, where it is stabilized by interactions involving N179 in A2 and R184 and S266 in A3 (Figure 3A, left). Following proteolytic cleavage, the CD shifts toward the A1 and A4 domains, weakening its contacts with the A2 and A3 domains. Thus, the CD moves out of this cavity, accompanied by stabilization of the C265–V271 segment in a new conformation (Figure 3A, right). The CD relocates closer to the A4 and A1 domains, forming contacts not only with the newly generated N-terminal segment but also with S275 in A4 and S86 and S88 in A1.

**Figure 3.**
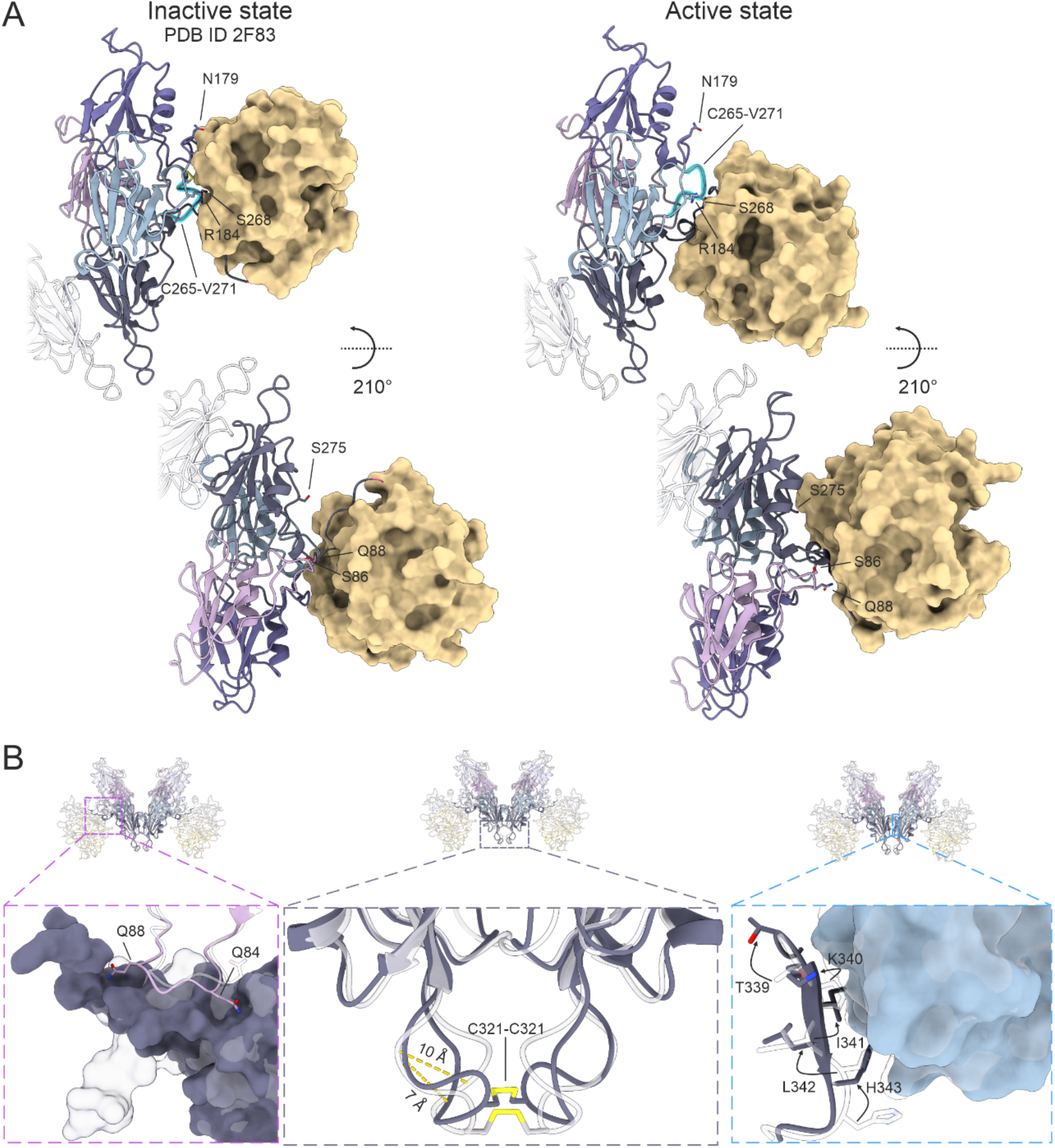
Structural rearrangements in AD disks upon activation of FXI. (A) Inactive FXI (zymogen), PDB ID: 2F83^3^, left, and FXIa, right, shown from two orientations. The first view highlights changes in the interaction of the CD with the A2 and A3 domains. The A3 segment undergoing a substantial conformational rearrangement is shown in cyan. The second view, after a 210° rotation, shows changes in the interaction of the CD with the A4 and A1 domains. (B) Interactions of the A4 domain (dark-violet) with other ADs in the inactive state (transparent grey) and active state (colored). Left: interaction of the A4 surface with the Q84-Q88 A1 loop (pink). Middle: interaction between the A4 domains near the connecting disulfide bond; the distance is measured between A319 and G324 within the same A4 domain. Right: interaction of the T339-H343 β-strand with the A3 domain surface (blue); arrows indicate reorganization of the β-strand amino acids’ side chains.

Activation-associated rearrangements are observed not only within the CD and at CD–apple-domain interfaces, but also in the interactions between the ADs themselves (Figure 3B). In the A1 domain, the side chains of Q84 and Q88 reorient toward A4, establishing a closer interdomain contact (Figure 3B, left). In addition, reorientation of the T339–H343 β-strand promotes closer interaction with the A3 domain (Figure 3B, right). Although the overall A4-A4 dimer interface remains similar in FXI and FXIa, local rearrangements are apparent near the intersubunit disulfide bond. These changes appear to permit closer intersubunit contacts, including potential polar interactions, that are less evident in the FXI zymogen. These structural changes seem to be facilitated by flexibility at the interface between the two monomers (Figure 3B, middle). According to the cross-species evolutionary conservation analysis of FXI, the discussed amino acids have average to high conservation score (Supplementary Figure 4).

### FXIa CD flexibility reveals additional active conformations and enzyme heterogeneity

While working on the C2 reconstruction of the FXIa homodimer (Figure 1B), we observed lower local resolution for the CDs (Supplementary Figure 1g). Upon relaxing the symmetry to C1, we found that the density corresponding to only one of the CDs in the FXIa dimer was markedly weaker, suggesting significant conformational flexibility. To further investigate this heterogeneity, the dataset was subjected to extensive 3D classifications, yielding three distinct states (Figure 4A, B). State 1 (37% of particles) resembles the C2 symmetric arrangement described in the previous sections, with both CDs adopting the same conformation, hereafter referred to as conformation ‘A’. In State 2 (27% of particles), one CD remains in conformation ‘A’ while the other adopts a distinct conformation ‘B’, positioned closer to the A1 domain. In State 3 (37 % of particles), one CD remains in conformation ‘A’, but the flexible CD is even further displaced towards A1 and adopts conformation ‘C’. Notably, the continuous displacement of one of the two CDs towards A1, from conformer ‘A’ to ‘C’, results in a gradual widening of the angle between A3/A4 and the CD, thereby further exposing the IRD motif (Figure 4C). The most optimal docking of the Factor IX Gla domain was observed for conformer ‘A’ (Figure 2C).

**Figure 4.**
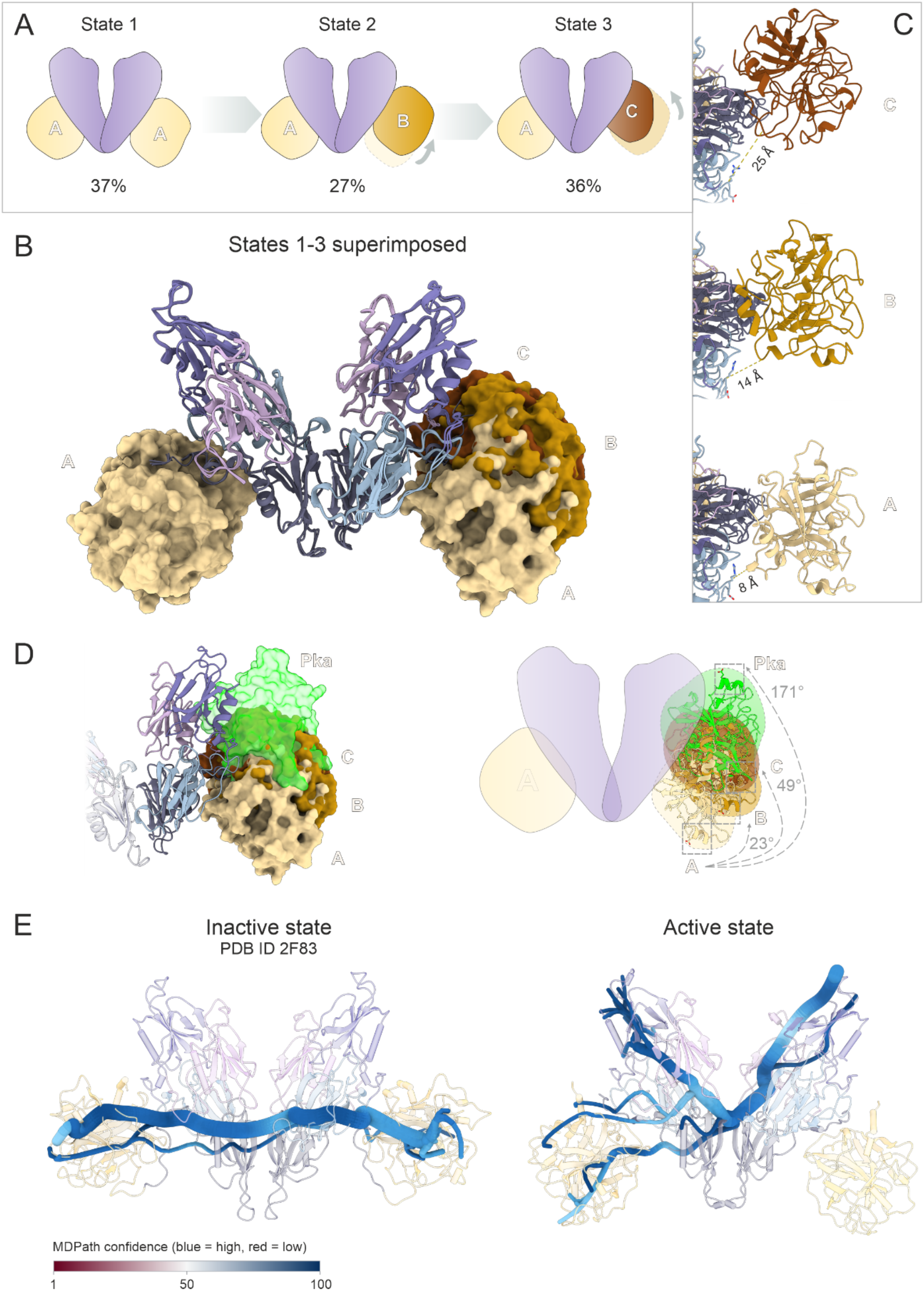
FXIa CD flexibility. (A) Schematic representation of the three conformational states of FXIa, with particle distributions indicated as percentages: State 1, CD conformation ‘A–A’, contains 37% of particles; State 2, CD conformation ‘A–B’, contains 27% of particles; State 3, CD conformation ‘A–C’, contains 36% of particles. The AD disc is colored violet; the CD in conformation ‘A’ is colored beige; the CD in conformation ‘B’ is colored light brown; and the CD in conformation ‘C’ is colored dark brown. (B) Superimposition of States 1–3. The CD surfaces in conformations ‘A’, ‘B’, and ‘C’ are colored as described in panel A. (C) Distance between the ^183^IRD^185^ motif and the CD in the different conformations is shown. The CD chains in conformations ‘A’, ‘B’, and ‘C’ are colored as described in panel A. (D) Analysis of CD orientation in the three FXIa states and PKa. The structures were compared following alignment of their AD platforms. The surface and ribbon representations of the FXIa CDs in conformations ‘A’, ‘B’, and ‘C’ are coloured as in panel A, while the PKa CD (PDB ID: 9VWT) is shown in light green. Relative CD rotations were measured using the Cα atom of N360 in FXIa State 1 as a common vertex. Vectors were defined from this vertex to the Cα atom of E525 in each FXIa state and to the corresponding E166 residue in PKa. (E) Allosteric communication calculated using MDPath, based on the mutual information of dihedral angles across all unbiased replicas for both the active and inactive states. Both calculations are highly confident, as indicated by the blue color, and are well sampled. Spline thickness correlates with the occurrence of the allosteric edge within the top 500 paths.

To further assess CD mobility within apple-domain-containing contact-system proteases, we compared the three apo-FXIa conformations (States 1–3) with the homologous PKa structure^4,48,51^ (Figure 4D, left). To quantify differences in CD orientation among the three FXIa states and relative to PKa, rigid-body rotations were analyzed. The AD platforms were first aligned, yielding an RMSD of 1.607 Å. The angular displacement of the conserved E525-containing region in the three FXIa states and the corresponding E166-containing region in PKa was then calculated using the align command in UCSF ChimeraX. The calculations were based on the Euler–Rodrigues rotation formula and the Kabsch algorithm^52,53^. Relative to FXIa State 1, PKa exhibited a pronounced rotation of 171°, whereas FXIa States 3 and 2 showed smaller, more localized screw-like rotations of 49° and 23°, respectively (Figure 4D, right). These findings demonstrate that the conserved AD architecture can accommodate markedly different CD orientations and interdomain interaction networks, which may contribute to substrate specialization. In FXIa, State 1 exposes and reshapes the A3/A4 surface implicated in FIX recognition.

It is intriguing that the homodimer exhibits asymmetric flexibility for one of the two CDs. To better understand this, the models of inactive and active states of FXI were analyzed using MDPath^47^. This analysis revealed that activation reshapes the way signals travel through the FXI dimer, with inactive FXI and FXIa states having markedly different allosteric communication networks (Figure 4E). In the inactive state, the two CDs are directly coupled, relaying allosteric signals across the dimer. In contrast, in the active state, one CD becomes fully uncoupled from the network, which may well explain the asymmetric heterogeneity and existence of additional active states. This rewiring upon activation is accompanied by a pronounced reorganization of the ADs: the dimer interface is extensively remodeled, driving the formation of two symmetric E286–R37 salt bridges. Each bridge directly links apple domains A1 and A4 at the core of the interface and propagates from there into the CD (Supplementary Figure 5).

## Discussion

FXI is a structurally distinctive coagulation protease and an important emerging antithrombotic target.^7^ It circulates as a disulfide-linked homodimer, with each subunit containing four ADs and a C-terminal CD.^1^ Despite previous structures of zymogen FXI^3,4^ and ligand-bound FXIa^36^, the structural consequences of activation and the organization of intact apo-FXIa have remained incompletely understood. Here, cryo-EM reveals extensive activation-associated rearrangements, a potential structural basis for FIX recognition, and pronounced and unexpected conformational heterogeneity in the FXIa CD.

Cleavage between R369 and I370 induces changes that extend far beyond the activation loop. The cleaved D359–R369 segment is repositioned, while the CD rotates and shifts relative to the AD platform. These movements reorganize CD-AD contacts and are accompanied by changes within the AD disc, including local rearrangements at the A4-A4 dimer interface. Thus, FXI activation involves global reorganization of the intact dimer rather than only local changes at the catalytic site.

Our MD analysis reveals that activation of FXI significantly alters signal transduction throughout the protein. In the inactive state, the two CDs are coupled, allowing allosteric signals to be passed back and forth. Upon activation, however, one of the CDs becomes disconnected from the network. This decoupling may account for the additional active states detected for one of the CDs in the cryo-EM structures of the homodimer.

We observe that as the CD transitions from conformer ‘A’ to ‘C’, it gradually exposes the previously occluded I183–R184–D185 region on A3, which has been implicated in FIX activation^6^. Successful docking of the FIX Gla domain into conformer ‘A’ (State 1) suggests that activation and the subsequent adjustment of the AD/CD angle create a sterically accessible A3/A4-facing surface capable of accommodating FIX, a site that remains hindered in the zymogen. This model is consistent with the established role of the A3 exosite and the sequential cleavage of FIX.^50,54^

Crucially, the presence of additional apo-FXIa conformations and pronounced mobility of the CD suggests that activation does not result in a single rigid active state but rather a dynamic conformational ensemble. The ability of the CD to occupy several positions relative to the AD platform, coupled with the variable opening of the CD/AD angle, suggests a mechanism of “dynamic scanning”. This plasticity likely allows the catalytic site to reposition itself relative to different AD exosites, facilitating the recognition and precise orientation of various macromolecular substrates.^8,55,56^

A comparison with PKa is hereby particularly informative, as PKa is the closest structural comparator for FXIa^4,57^. FXI arose through duplication of the prekallikrein gene, which explains their shared four-apple-domain architecture. FXI and PKa are closely related evolutionarily and functionally within the contact system, and both circulate in complex with high-molecular-weight kininogen, a physiological substrate of PKa.^8,58^ Comparison with the homologous PKa full-length crystal structure reveals that, within this conserved domain architecture^4^, the structurally similar AD platform displays the CD in a fourth orientation, distinct from conformers ‘A’, ‘B’, and ‘C’ of FXIa. While the available full-length PKa crystal structure provides only a static snapshot and may not fully capture the extent of CD mobility^4,59^, the existence of distinct CD conformers in FXIa and PKa suggests that CD plasticity might be a shared feature of apple-domain-containing contact-system proteases. This plasticity likely enables the catalytic site to dynamically adapt to the requirements of different substrates.

Although the proposed FIX-binding mode and the functions of the different FXIa conformations require further validation, our structures show that FXI activation reorganizes the entire dimer into a flexible, multi-state enzyme. This dynamic architecture provides several novel targets for inhibition beyond the catalytic site, including the exposed A3/A4 surface, the CD-AD interfaces, and the regions controlling CD movement. By stabilizing an inactive conformation or limiting CD mobility, small molecules or antibodies could inhibit FXIa through a non-catalytic mechanism, providing a path toward more specific antithrombotic therapies.

## Supporting information

Supplementary Video 1_Transition from the FXI zymogen to the activated FXIa state

Supplementary Video 2_Conformational variability of activated FXI

## Associated content

### Supporting Information

The Supporting Information is available free of charge and includes the following:

**Supporting Information:** Supplementary Methods, Figures S1–S5, and Tables S1–S2 (PDF).

**Supplementary Video 1:** Transition from the FXI zymogen to the activated FXIa state, shown from bottom and top views (MP4).

**Supplementary Video 2:** Conformational variability of activated FXI. Morphs between the FXIa cryo-EM 3D classes shown in Figure 4A,B (MP4).

## Acknowledgements

We thank George Broutzakis for his assistance during the initial stages of data processing. We are also grateful to Dr. Anja Blanque for recording the second cryo-EM dataset. The authors acknowledge the funding support received from the German Research Foundation (DFG) (KA 5558/1-1).

