## Supplementary Information for "Cryo-EM structures of apo human Factor XIa reveal catalytic-domain flexibility and exposure of the Factor IX-binding site"

**Contents**

| | **Page** |
|---|---|
| Supplementary Figure 1: Negative stain and cryo-EM analysis of FXIa | S2 |
| Supplementary Figure 2: Interactions between CD and the D359–R369 fragment in inactive and active states of FXIa. | S3 |
| Supplementary Figure 3: Proteolysis-driven structural and electrostatic activation of FXI. | S3 |
| Supplementary Figure 4: Cross-species evolutionary conservation analysis of FXI. | S4 |
| Supplementary Figure 5: Allosteric connectivity of the dimer. | S4 |
| Supplementary Table 1: Cryo-EM data collection, refinement and validation statistics | S5 |
| Supplementary Table 2 | S6 |
| Supplementary methods | S7 |
| References | S12 |

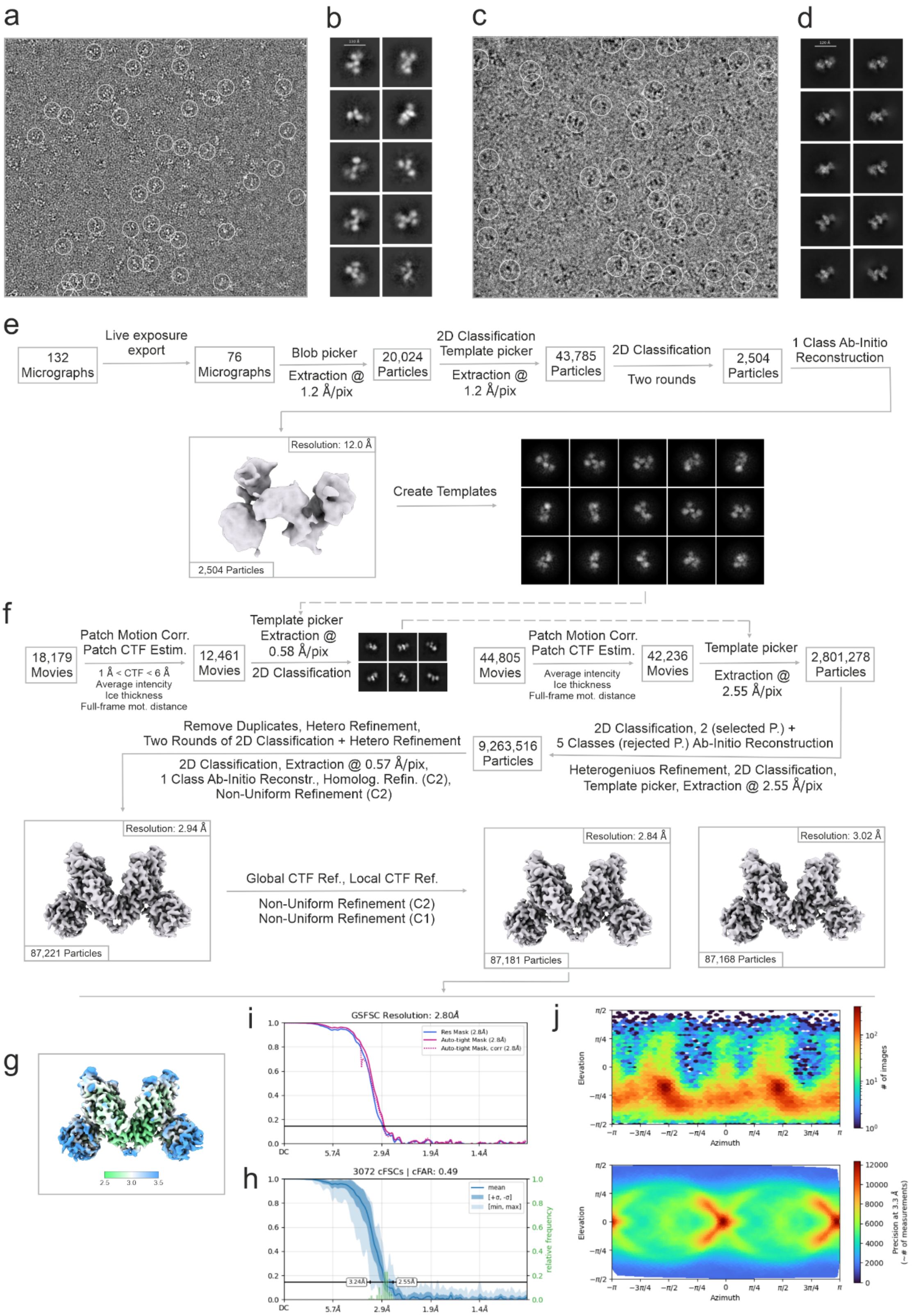


**Supplementary Figure 1: Negative stain and cryo-EM analysis of FXIa.** (a) Representative NS micrograph. (b) Representative NS 2D class averages. (c) Representative cryo-EM micrograph (d) Representative cryo-EM 2D class averages. (e) Flowchart for NS data processing. (f) Flowchart for cryo-EM data processing. (g) Final cryo-EM volume colored according to the local resolution. (h) Gold standard FSC. (i) Summary plot of the 3,072 conical Fourier Shell Correlation (cFSC) curves. The global map anisotropy is quantified by the Conical FSC Area Ratio. (j) Euler angle distribution.

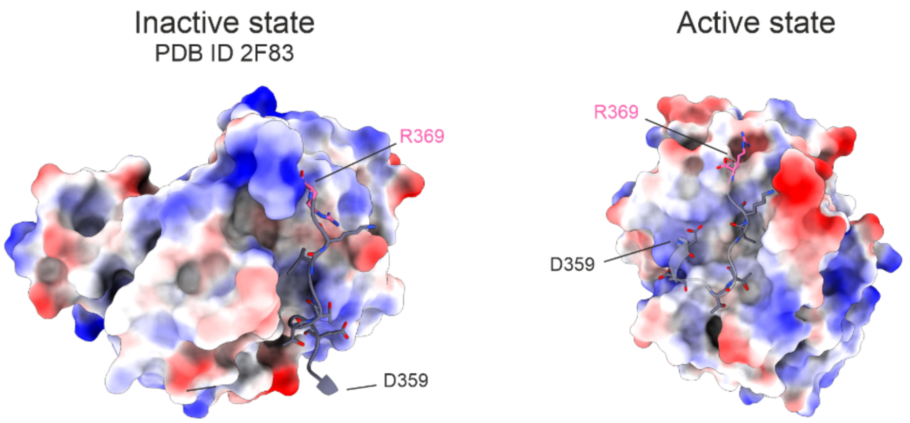


**Supplementary Figure 2: Interactions between CD and the D359–R369 fragment in inactive and active states of FXIa.**

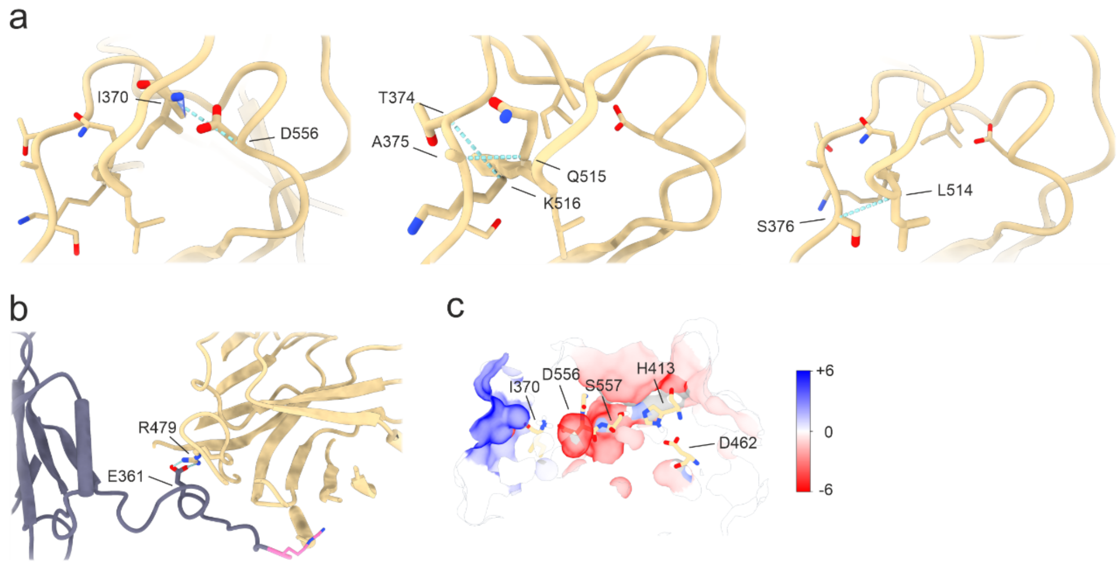


**Supplementary Figure 3: Proteolysis-driven structural and electrostatic activation of FXI.** (a) Schematic representation of the three-step FXI activation mechanism. Proteolytic cleavage triggers insertion of the newly formed I370 N-terminus and formation of the I370–D556 salt bridge (left), followed by slow repacking of the conserved protease-domain core through contacts such as A375–Q515 (middle); the final active conformation is further stabilized by additional hydrogen bonds, such as S376–L514 (right). (B) Close-up of the newly formed C-terminus (pink) after cleavage, which is held in position by the rapid formation of an E361–R479 salt bridge. (C) Overlay of the electrostatic potential of the active-state catalytic domain, comparing the pre- and post-salt-bridge states, computed using QM/MM calculations. Salt-bridge formation decreases the electrostatic potential at the catalytic triad, indicating stabilization of the positively charged catalytic histidine formed during catalysis and thereby promoting electrostatic preorganization of the active site. The electrostatic-potential difference is shown using a colour scale ranging from −6 kcal $mol^{-1}$ $e^{-1}$ (deep red) to +6 kcal $mol^{-1}$ $e^{-1}$ (deep blue).

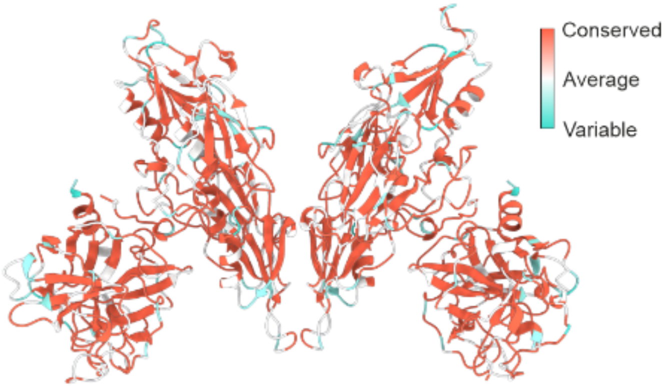


**Supplementary Figure 4: Cross-species evolutionary conservation analysis of FXI.** The Cryo-EM structure of human FXIa (current study) is displayed as a ribbon model. Amino acid conservation scores were mapped onto the structural coordinates following a multiple sequence alignment (MSA) of cross-species orthologs including Human (Homo sapiens), Mouse (Mus musculus), Bovine (Bos taurus), and Pig (Sus scrofa). Conservation values were calculated using the AL2CO numerical entropy algorithm in UCSF ChimeraX. Residues are colored along a continuous gradient from highly variable (turquoise; minimum score: -2.81) through average baseline conservation (white; midpoint score: -1.07) to hyper-conserved regions (tomato; maximum score: 0.67).

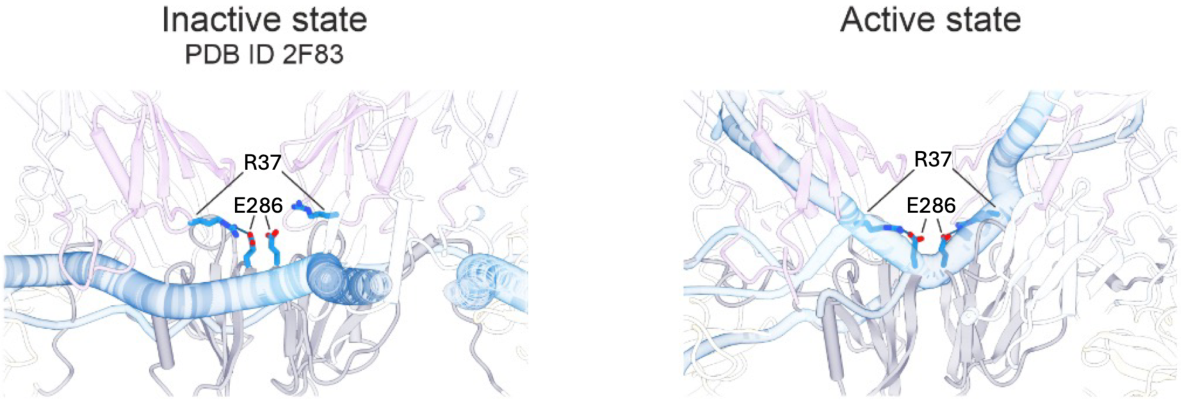


**Supplementary Figure 5: Allosteric connectivity of the dimer.** A close-up of Figure 4E. Both dimer interfaces display allosteric communication (blue tube). The residues directly involved in the connection are highlighted as blue sticks. Salt bridges involving R37 and E286 facilitate allosteric communication to the AD in the active state, whereas those paths are absent in the inactive state.

**Supplementary Table 1: Cryo-EM data collection, refinement and validation statistics.**

| | |
|---|---|
| **Data collection and processing** | |
| Microscope | Titan Krios G4 (Selectris X, E-CFEG) |
| Magnification | 270,000 |
| Voltage (kV) | 300 |
| Camera | Falcon 4i |
| Pixel size (Å) | 0.571 |
| Total electron dose ($e^-$/Å$^2$) | 50 |
| Defocus range (µm) | -0.8 – -2.8 |
| Movies (No.) | 44,805 |
| Number of frames | 50 |
| Initial particle images (No.) | 9,263,515 |
| Final particle images (No.) | 87,181 |
| Symmetry imposed | C2 |
| Map resolution (Å) | 2.80 |
| Map resolution range(Å) | 2.55-3.24 |
| FSC threshold | 0.143 |
| **Atomic model composition** | |
| Chains | 4 |
| Symmetry imposed | C2 |
| Non-hydrogen (protein) atoms | 9,442 |
| Residues | 1,214 |
| Ligands | 10 (NAG) |
| **Refinement (Phenix)** | |
| RMSD bond lengths (Å) | 0.003 (0) |
| RMSD angles (°) | 0.598 (0) |
| Model to map fit, CC mask | 0.84 |
| Model to map fit, CC box | 0.79 |
| B-factor (mean, Å$^2$)<br>Protein<br>Ligands | <br>72.88<br>157.06 |
| **Validation** | |
| Clashscore | 2.36 |
| Ramachandran outliers (%) | 0.00 |
| Ramachandran favoured (%) | 97.35 |
| Ramachandran allowed (%) | 2.65 |
| Molprobity score | 1.14 |
| EMRinger score | 5.44 |

**Supplementary Table 2:** Input features comprising the exported Deep-TDA CV: 41 features (74 columns) retained by feature-level unsupervised filtering from 48 differentiable candidates (86 columns). The full specification defines 55 physical descriptors (100 columns); seven alignment-dependent rotation descriptors (#11–17; 14 columns) were excluded before model construction. Feature IDs span 1–92 but are not a feature count or output-column indices; #39–40 are intentionally unused and #48–82 are reserved (see note c). Residue numbers are in mature UniProt P03951 numbering (Ile370 is the new N-terminus produced by Arg369–Ile370 cleavage; His413/Asp462/Ser557 are the catalytic triad; Asp556 is the activation salt-bridge partner). Per-monomer features are evaluated for both monomers (A and B) and contribute two columns each; the inter-subunit group is global. The use of physically interpretable structural descriptors as inputs to a Deep-TDA model follows established data-driven-CV practice.[1-3]

| Feature group | # feat.[a] | Definition (residues, mature P03951; candidate index) |
|---|---|---|
| Protease-domain orientation | 9 | Centre-of-mass distance of the protease domain (388–625) to the N-terminal/apple domains 1–90, 91–180, 181–270, 271–360 (#1–4); Cα distances 388–184, 555–184, 555–263, 480–320, 500–340 (#5, #7–10). FXI architectural rationale.[4,5] |
| Activation latch | 12 | Cα distances 362–270, 365–268, 367–178, 370–360 (#18–20, #29); backbone φ, ψ of residue 365 and φ of residue 369, each as sin/cos (#21–26); coordination numbers of the activation loop (360–370) with 260–270 and with 320–360 ($r_0 = 6.5$ Å, $n = 6$, $m = 12$; #27–28). Activation-domain rationale[6-9]; rational coordination-function convention.[10] |
| Salt bridge and catalytic triad | 6 | Ile370 N to the Asp556 carboxylate: distance to the Oδ1/Oδ2 midpoint (#30) and to Oδ1 (#31); catalytic-triad contacts His413 Nε2–Asp462 Oδ1 (#33) and Ser557 Oγ–His413 Nε2 (#34); backbone N555–N557 (#37); Ile370 ψ dihedral (#41). Canonical chymotrypsin-family activation-switch and active-site rationale[6-9,11], with the FXI-specific Ile370–Asp556 assignment supported by.[12] |
| Protease-domain rotation | 6 | Cα distances 557–45, 557–225, 413–315, 413–135, 390–315, 604–45 (#42–47). FXI protease/apple-domain orientation rationale.[4,5] |
| Inter-subunit asymmetry[b] | 8 | Inter-subunit Cα distance (residue 321; #84), inter-subunit coordination (271–360; #85) and centre-of-mass distance (181–270; #92); A−B difference ("delta") and A/B dispersion ("sigma") of orientation #1 (#86, #89), salt bridge #30 (#87, #90) and latch coordination #27 (#88). FXI dimer and transactivation rationale.[4,5] |

[a] Number of features per monomer for the first four groups, each contributing two columns (A and B).
[b] Global inter-subunit group (8 columns). Totals: 33 × 2 + 8 = 74 columns (41 features).
[c] Feature IDs #39 and #40 were deliberately left unused: they would have encoded sin φ(370) and cos φ(370), but φ(370) spans the Arg369–Ile370 scissile bond and is therefore a genuine backbone torsion only in the uncleaved zymogen; after cleavage it becomes a cross-chain pseudo-torsion. ψ(370) (#41) was retained because its defining atoms remain on the FXIa light chain. IDs #48–82 are reserved and unassigned. The excluded alignment-dependent descriptors #11–17 comprised quaternion/rotation, PCAVARS, and dRMSD terms.[13-15]

### Supplementary methods

#### QM/MM electronic-structure calculations

The electronic contribution of the activation salt bridge was evaluated using an electrostatically embedded QM-cluster/QM/MM model of the activated FXIa catalytic domain at a fixed cleaved-state geometry, implemented in Psi4 1.9.1.[16] Two electronic states were calculated on the same heavy-atom structure: the formed ion pair (Ile370 $^{+}NH_3$/Asp556 $COO^{-}$; N···O = 2.75 Å) and the broken pair (neutral $NH_2$/COOH). Because the two residues are collectively neutral in both states, the QM region retained the same total charge and multiplicity (−1, singlet). Accordingly, the density difference, $\Delta\rho = \rho_{\mathrm{formed}} - \rho_{\mathrm{broken}}$, isolates the effect of ion-pair charge formation independently of activation-associated conformational changes.

The 202-atom QM region comprised 22 residues, including the catalytic triad (His413, Asp462, and Ser557), the oxyanion-hole amides (Gly555 and Ser557), the Ile370–Asp556 pair, and all side chains within 3.5 Å. Boundaries were introduced only across non-polar Cα–Cβ or backbone Cα–C/N–Cα bonds and capped with hydrogen link atoms, preserving all amide and disulfide bonds. The remaining protein atoms, neutralizing ions, and active-site waters were represented by 19,474 fixed CHARMM36m point charges,[17] identical in both states, with charge redistribution applied at each QM/MM boundary. Bulk solvent was described using CPCM with $\varepsilon = 78.39$.[18] No additional protein-interior dielectric was applied because the protein environment was represented explicitly.

Single-point energies, electron densities, and electrostatic potentials were calculated at the ωB97X-D/def2-TZVP level[19,20] using density-fitted SCF[21], the def2-universal-jkfit auxiliary basis[22], and an SCF convergence threshold of $10^{-8}$. Differences in electron density and electrostatic potential, atomic populations, and non-covalent interaction descriptors[23] were evaluated on a common grid. The interaction energy was benchmarked using a methylammonium–acetate model. The ωB97X-D value (−126.0 kcal $mol^{-1}$) agreed within approximately 1.5 kcal $mol^{-1}$ with counterpoise-corrected DF-MP2/def2-TZVP (−124.5 kcal $mol^{-1}$) and SAPT0 (−124.8 kcal $mol^{-1}$) and was converged relative to CP-MP2/def2-QZVP (−124.5 kcal $mol^{-1}$).

#### DeepCV constructions and enhanced sampling

Training data for the collective variable (CV) comprised labelled frames from unbiased simulations of the FXI zymogen and activated FXIa, sampled every 100 ps. Because the two systems differ in chain composition, residues were mapped to the common numbering of UniProt P03951 by pairwise sequence alignment using Biopython.[24] Features were defined in this numbering and resolved independently for each topology; explicit atom indices were taken from the active-state build, whose ordering matches the production topology.

Each frame was featurised with MDAnalysis[25,26] using descriptors of protease-domain orientation, the activation latch, the Ile370–Asp556 salt bridge and catalytic-triad geometry, protease-domain rotation, and inter-subunit asymmetry. Per-monomer descriptors were calculated for both monomers and were additionally represented by inter-monomer difference and dispersion terms. The initial specification comprised 55 descriptors represented by 100 columns: 45 per-protomer descriptors were evaluated for both monomers (90 columns), and 10 inter-subunit descriptors contributed one column each. The seven structural-alignment-dependent rotation descriptors (#11–17) were emitted as NaN placeholders and excluded before differentiable model construction, leaving 48 candidate features represented by 86 columns. Feature-level unsupervised filtering then removed near-constant features (relative standard deviation below 0.05) and redundant features (absolute Pearson correlation above 0.95). The final exported Deep-TDA input comprised 41 features represented by 74 columns and evaluated from 1,208 atoms.

A one-dimensional CV was trained using the Deep Targeted Discriminant Analysis objective implemented in mlcolvar[2,3] with a PyTorch backend.[27] The two states were assigned Gaussian targets centred at −1 and +1 with widths of 0.5, using the default loss weights $\alpha = 1$ and $\beta = 100$. Generalisation was assessed by five-fold leave-one-replica-out cross-validation, with one of the five replicas from each state withheld in each fold. Input features were z-scored using statistics calculated exclusively from the corresponding training replicas.

Network architecture and optimisation parameters were selected with Optuna[28] using a Tree-structured Parzen estimator and median pruning over 40 trials. The search included one or two hidden layers, widths of 16, 32, or 64 neurons, learning rates from $1 \times 10^{-4}$ to $1.5 \times 10^{-3}$, batch sizes of 128, 256, or 512, and $L_2$ weight decay from $1 \times 10^{-6}$ to $1 \times 10^{-2}$. Trials minimised the mean cross-validation loss and used early stopping on the held-out replicas. The deployed model was refitted using full-dataset standardisation statistics and early-stopped against a held-out replica. Among several random seeds, the model with the lowest held-out loss was retained; it contained one hidden layer of 16 neurons and a weight decay of $2.2 \times 10^{-6}$. Feature calculation, standardisation, and the trained network were combined into a single module mapping the selected Cartesian coordinates directly to the scalar CV and exported as a traced TorchScript model.

Biased simulations were performed with OpenMM 8.4[29] and the GLUED GPU-accelerated Library for Unified Exploration Dynamics plugin ([https://github.com/MarvinTaterra/GluedMD/tree/7cec9a5aaf8d5f3b2484edeff746f2a18e7b593e]). GLUED evaluates the TorchScript CV on the GPU and obtains biasing forces by automatic differentiation at every integration step. A two-dimensional bias was applied to the Deep-TDA CV and the monomer-A Ile370(N)–Asp556 carboxylate distance, corresponding to feature 30, using the well-tempered explore variant of On-the-fly Probability Enhanced Sampling (OPES).[30,31] The bias factor was $\gamma = 20$, corresponding to approximately 50 kJ mol$^{-1}$, and kernels were deposited every 500 steps (1 ps). Kernel widths were set to approximately half the within-basin standard deviation of the unbiased active ensemble: $\sigma_{\mathrm{CV}} \approx 0.25$ and $\sigma_{\mathrm{salt\ bridge}} \approx 0.013$ nm. Soft harmonic walls restricted sampling to CV values between −2 and 2 and salt-bridge distances between 0.2 and 2.8 nm.

Sampling was initiated from the equilibrated active state. Ten independent single-GPU walkers, corresponding to five active-state replicas with two random seeds each, were started from the same restart coordinates with independently assigned Maxwell–Boltzmann velocities and thermostat and barostat seeds. Each walker was propagated for 100 ns. Simulation settings otherwise matched the unbiased simulations, including PME electrostatics, force-switched van der Waals interactions, NPT conditions at 303.15 K, a 2 fs integration step, and constraints on bonds involving hydrogen.

The two CVs, total applied bias (V(s)), and OPES normalisation term (c(t)) were recorded every 20 ps. Frame weights were calculated as $w = \exp(\frac{V(s)-c(t)}{k_{\mathrm{B}}T})$. Coordinates of the feature atoms were stored for subsequent DeepTICA analysis.[32]

#### Slow-mode analysis of the enhanced-sampling trajectories

Each frame of the retained OPES-Explore ensemble was represented by 293 descriptors, corresponding to 576 protomer-resolved columns. The feature set comprised 69 salt bridges and 50 coevolution-derived residue pairs, encoded using rational switching functions of the minimum charged-atom or Cα distance ($r_0 \approx 4.5$ Å) to provide bounded formed/broken values between 0 and 1; 50 graded Cα distances selected by a Δ-contact screen of the equilibrium end-state ensembles; 76 backbone φ/ψ torsions from the activation region, represented by their sine and cosine; and the engineered descriptors of protease-domain orientation, the 361–387 activation latch, the active site, and the inter-subunit interface defined above. Residues were indexed using FXI numbering (UniProt P03951) and mapped independently onto each topology by global sequence alignment. Features were extracted with MDAnalysis.[25,26]

Coevolutionary couplings were obtained by querying the FXI protease domain, residues 370–607, against the ColabFold MMseqs2 server.[33] The resulting alignment contained 8,478 sequences, of which 4,702 remained effective after query anchoring and gap filtering. Couplings were inferred by pseudo-likelihood maximisation direct-coupling analysis with average-product correction using plmDCA as implemented in pydca.[34,35] Motivated by previous work showing that coevolving residue pairs absent from a reference structure can report alternative functional conformations[36], the 50 highest-scoring FXI pairs were included as candidate state-dependent contacts and annotated by their Cα distances in both end states.

Because the trajectories were biased, each frame was assigned the OPES reweighting factor $w = \exp(\frac{V_{\text{bias}} - c(t)}{k_B T})$.
Time-lagged configuration pairs were constructed only within individual walkers, never across walkers, using a lag time of 200 ps, corresponding to ten saved frames. Slow collective modes were identified using both time-lagged independent component analysis (TICA)[37,38] and DeepTICA.[32] Both methods were applied to the same reweighted feature set and lag time using mlcolvar 1.3.1[3] with a PyTorch 2.10 backend.[27] DeepTICA networks were trained with an $L_2$ weight decay of $1 \times 10^{-4}$, early stopping with a patience of 20 epochs, and three random seeds. Generalisation was evaluated by withholding one complete walker, rather than randomly splitting frames, to prevent leakage between temporally correlated configurations.

The leading slow component satisfied three validation criteria: its generalised eigenvalue was within the physically admissible interval $\lambda_1 = 0.86$; its implied timescale plateaued at approximately 200 ps for lag times of 200–400 ps on the held-out walker; and the difference between training and held-out autocorrelations was approximately 0.06, below the predefined overfitting threshold of 0.12. This gap was unaffected by a tenfold increase in weight decay, indicating that it primarily reflected between-walker variability rather than model over-parameterisation. Higher components were excluded because their eigenvalues fell outside $[-1,1]$. This instability was attributed to the low effective sample size of the reweighted ensemble (ESS $\approx 550$ of 50,000 frames), which rendered the weighted covariance matrices ill-conditioned for higher generalised-eigenvalue modes.

Feature contributions to the leading component were quantified by Integrated Gradients[39] over all 50,000 frames. Reduced models were then trained using the top 40, 80, 120, or 160 ranked features. The 120-feature model retained approximately 86% of the held-out implied timescale of the full model (206 versus 240 ps) while using fivefold fewer, directly interpretable descriptors. It was therefore used for mechanistic interpretation, whereas the full model was retained for analyses requiring the longest recovered timescale.

Individual interactions were positioned along the transition using two complementary measures. Structural differences between the equilibrium end states were quantified by *Δ*Form, defined as the difference in interaction occupancy between the active and zymogen ensembles. Their transition behaviour was characterised by correlation with $cv_0$, occupancy profiles binned along $cv_0$, the $cv_0$ value at which occupancy crossed 0.5, and the magnitude of their loading or attribution on the slowest TICA or DeepTICA component.

For mechanistic ordering, cv0 was oriented from zymogen-like toward active-like configurations. Because all enhanced-sampling trajectories contained cleaved FXIa, proteolysis itself was not simulated. Cleavage-dependent interactions, including Ile370 insertion and formation of the Ile370–Asp556 ion pair, were therefore placed at the beginning of the mechanism from their equilibrium end-state ΔForm values, not from an observed formation time. The ordering of subsequent post-cleavage rearrangements was inferred from cv0-binned occupancy profiles together with their TICA/DeepTICA loadings or attributions. Implied timescales were used for lag-time convergence and model comparison and were not interpreted as experimental activation rate constants.

**Electronic effect of the Ile370–Asp556 salt bridge**

The Ile370–Asp556 ion pair is generally considered a structural latch that anchors the liberated N-terminus in the activation pocket and stabilizes the final protease conformation[12,40] To isolate its electronic contribution, we performed QM/MM calculations on two states with identical heavy-atom geometries, point-charge environments, and total QM-region charge. Thus, all differences arise solely from the ion-pair charge state rather than activation-associated conformational changes.

Ion-pair formation redistributes 1.37 e of electron density and alters the electrostatic potential across the catalytic triad. Although Asp556 is 7.8 Å from the Ser557 Oγ nucleophile and does not contact it directly, the potential becomes more negative by −7.6 kcal $mol^{-1}$ $e^{-1}$ at Ser557 Oγ, −4.5 at His413 Nε2, and −3.1 at Asp462 Oδ1. The distance-dependent decay and absence of a continuous interaction pathway in the NCI analysis support a through-space electrostatic effect rather than covalent or hydrogen-bond-mediated coupling.

This negative potential preferentially stabilizes the positively charged catalytic histidine formed after proton transfer from serine, which is present in the transition state and tetrahedral intermediate but not in the resting enzyme. Ion-pair formation may therefore lower the activation barrier by electrostatically preorganizing the catalytic site. A comparable long-range effect has been reported for subtilisin, where an outer-shell negative charge approximately 8 Å from the active site contributes more than 2 kcal $mol^{-1}$ to catalysis,[41] suggesting that such electrostatic preorganization may be a recurring feature of serine proteases.

**Mechanism of FXI Activation**

To determine how factor XI transitions from the inactive zymogen to the active protease FXIa following proteolytic cleavage, we combined equilibrium molecular dynamics simulations of both end states with enhanced sampling along a machine-learned collective variable. Formation of the canonical activation salt bridge between the liberated Ile370 N-terminus and Asp556, enabled by cleavage of the Arg369–Ile370 bond, provided a structural marker of activation. Repeated forward and reverse transitions indicated reversible sampling of the activation pathway.

We next trained a DeepTICA[32] model on the enhanced-sampling ensemble to identify the slowest collective motion underlying the transition. In contrast to the state-discriminating collective variable used for sampling, DeepTICA was optimized to recover the dominant slow mode and was interpreted alongside its linear TICA counterpart. Individual interactions were characterized using two complementary metrics: ΔForm, defined as the difference in occupancy between the active and zymogen states, and their behavior along the activation coordinate $cv_0$, including the timing and steepness of formation and their association with the slowest mode. Together, these analyses resolved a three-step activation cascade.

The first step is initiated directly by cleavage. The liberated Ile370 side chain inserts into a hydrophobic pocket of the protease domain, with the Ile370–residue 517 and Ile370–residue 498 contacts showing nearly complete formation upon activation (ΔForm = +0.89 and +0.86, respectively). Simultaneously, the newly exposed N-terminus forms the Ile370–Asp556 activation salt bridge, which is absent in the zymogen and present in 99% of active-state frames (ΔForm = +0.99). The cleaved terminus is further stabilized by the 366–600 contact (ΔForm = +0.73) and the 371–551 hydrogen bond (ΔForm = +0.80). Although these interactions represent the largest structural differences between the end states, they form directly after cleavage and therefore act as a trigger that primes the protease domain for subsequent reorganization.

The second step comprises a slower repacking of the protease-domain core around the docked N-terminus and represents the dominant slow conformational process. Its central event is formation of the coevolving 375–515 contact, whose occupancy increases from approximately 67% early in the transition to approximately 90% in the active state, with a midpoint near $cv_0 \approx 0.7$. This interaction makes the strongest contribution to the leading DeepTICA/TICA mode. Core consolidation is accompanied by formation of the 374–516 hydrogen bond (ΔForm = +0.57) and the 361–479 salt bridge (ΔForm = +0.47), with the latter forming near $cv_0 \approx 0.6$. The association of the 375–515 contact with both the slowest mode and the central region of the activation coordinate identifies this repacking event as the principal conformational bottleneck of activation. Its localization within an evolutionarily coupled network further suggests that the activation mechanism is encoded in the coevolutionary architecture of the protease domain.

In the final step, consolidation of the core is followed by formation of the 376–514 hydrogen bond (ΔForm = +0.57), which occurs late in the transition, near $cv_0 \approx 0.8$, and locks the protease domain into its mature active conformation. The coevolving network does not tighten uniformly during this process. While the 375–515 contact consolidates, other coupled pairs, including 517–550 and 551–588, transiently weaken and do not contribute directly to the forward activation cascade.

The minimal mechanistic model therefore comprises three principal events: Ile370 insertion and formation of the I370–D556 salt bridge as the cleavage-triggered switch, formation of the 375–515 coevolving-core contact as the dominant slow conformational step, and formation of the 376–514 hydrogen bond as the final lock-in of the active protease. The remaining electrostatic and hydrogen-bond changes either arise directly from cleavage or occur as secondary consequences of core consolidation.
